# Impact of Segmentation Errors in Analysis of Spatial Transcriptomics Data

**DOI:** 10.1101/2025.01.02.631135

**Authors:** Jonathan Mitchel, Teng Gao, Eli Cole, Viktor Petukhov, Peter V. Kharchenko

**Affiliations:** Department of Biomedical Informatics, Harvard Medical School, Boston, MA; Program in Health Sciences & Technology, Harvard Medical School & Massachusetts Institute of Technology, Boston, MA; San Diego Institute of Science, Altos Labs, San Diego, CA

**Author notes:** These authors contributed equally (Jonathan Mitchel, Teng Gao).

## Abstract

Spatial transcriptomics aims to elucidate cell coordination within biological tissues by linking the state of the cell with its local tissue microenvironment. Imaging-based assays are particularly promising for exploring such interdependencies, as they can resolve molecular and cellular features with subcellular resolution in three dimensions. Quantification and analysis of cellular state in such data, however, ultimately depends on the ability to recognize which molecules belong to each cell. Despite computational and experimental progress, this cell segmentation task remains challenging. Here we re-analyze data from multiple tissues and platforms and find that segmentation errors currently confound most downstream analysis of cellular state, including analysis of differential expression, inference of neighboring cell influence, and ligand-receptor interactions. The extent to which mis-segmented molecules impact the results can be striking, often dominating the set of top hits. We show that factorization of molecular neighborhoods can be effective at isolating such molecular admixtures and minimizing their impact on downstream analysis, analogous to doublet filtering of scRNA-seq data. As applications of spatial transcriptomics assays become more widespread, we expect corrections for the confounding effect of segmentation errors to become increasingly important for being able to resolve molecular mechanisms of tissue biology.

## Introduction

Spatial transcriptomics assays are designed to capture the transcriptional state of cells within their native tissue context^1^. The central promise of these rapidly advancing techniques lies in their potential to elucidate how cells coordinate and interact within their microenvironment, providing improved understanding of the mechanisms governing tissue organization and function^2^. In pursuit of these insights, a variety of computational methods have been developed, often drawing on approaches used in bulk or single-cell RNA sequencing analysis^3^. These include differential expression tests for comparing cellular states across distinct tissue microenvironments, analysis of general associations between cellular state and the properties of the immediate neighborhood of the cell, or identification of cognate ligand-receptor pairs that could mediate cellular interactions. These techniques are being actively applied in diverse biological settings, including tumor-immune interactions^4–11^, age-associated tissue distortions^12,13^, and developmental processes^14–16^.

There currently are two major categories of assays: sequencing- and imaging-based spatial transcriptomics. Sequencing-based methods, such as Slide-seqV2^14^, hybridize RNA on positionally-barcoded surfaces. Their ability to resolve the state of individual cells is limited by the resolution of the surface features, lateral diffusion, and more fundamentally by the ambiguity that arises when using a two-dimensional surface to map the three-dimensional cell arrangements which are present even in thin ∼10 micron slices. Consequently, computational analyses of such data generally avoid attribution of molecules to specific cells and try to mitigate the uncertainty of cellular state in other ways^17^.

In contrast, imaging-based methods, such as MERFISH^18^ or STARmap^19^, are in principle devoid of such limitations. They can detect individual RNA molecules with nanometer-scale spatial resolution, retaining subcellular depth information across three-dimensional volumes. However, answering questions about cellular state with such data depends on the ability to accurately segment the cell. Most segmentation methods rely on additional microscopy stains, including DAPI, polyA, and plasma membrane stains, to estimate the likely boundaries of each cell^20,21^. Segmentation can also be improved by considering the molecular composition of a cell^22,23^.

Despite these aids, accurate segmentation remains challenging due to uncertainties in staining signals, the presence of non-specific background molecules, and limited three-dimensional scope. Consequently, the transcriptional composition of the segmented cells can also be imprecise, with some molecules from neighboring, sometimes unobserved, cells being mistakenly attributed to a given cell.

In this study, we examine the impact of such segmentation errors on the downstream analysis of imaging-based spatial transcriptomics data. Re-analyzing data from different platforms, we demonstrate that for common analysis tasks, such context-dependent differential expression or ligand-receptor interactions, the artifacts of the segmentation errors dominate the results. Nevertheless, we show that low-dimensional decomposition of molecular neighborhoods can mitigate the impact of such admixtures on existing analysis methods.

## Results

### Analysis of cell type markers reveals admixtures from neighboring cells

To assess potential impact of segmentation errors, we examined several imaging-based spatial datasets from tissues where a matching scRNA-seq dataset was also available. These included MERFISH measurements of mouse hypothalamus^24^ and ileum^22^, CosMx data on non-small cell lung cancer (NSCLC)^25^, and Xenium data from an FFPE sample of human pancreas that utilizes the recent Multimodal Cell Segmentation kit (from 10x genomics website).

We first examined how cell type expression markers derived from the scRNA-seq data were expressed in the spatial data. Using cell segmentations from the original publications and manually matching cell type annotation names, we observed frequent presence of cell type marker transcripts in other cell types in spatial data (Figure 1a,b). Such non-specific expression was more prevalent for markers of certain cell types. For instance, in the NSCLC dataset, background expression of fibroblast markers, such as *COL3A1* or *COL1A2*, was noticeable in almost all other cell types (Figure 1a). Similarly, in mouse ileum data, markers of enterocyte population, such as *Slc5a1* and *Slc51a*, were frequently detected in most of the other cell types (Figure 1b).

**Figure 1.**
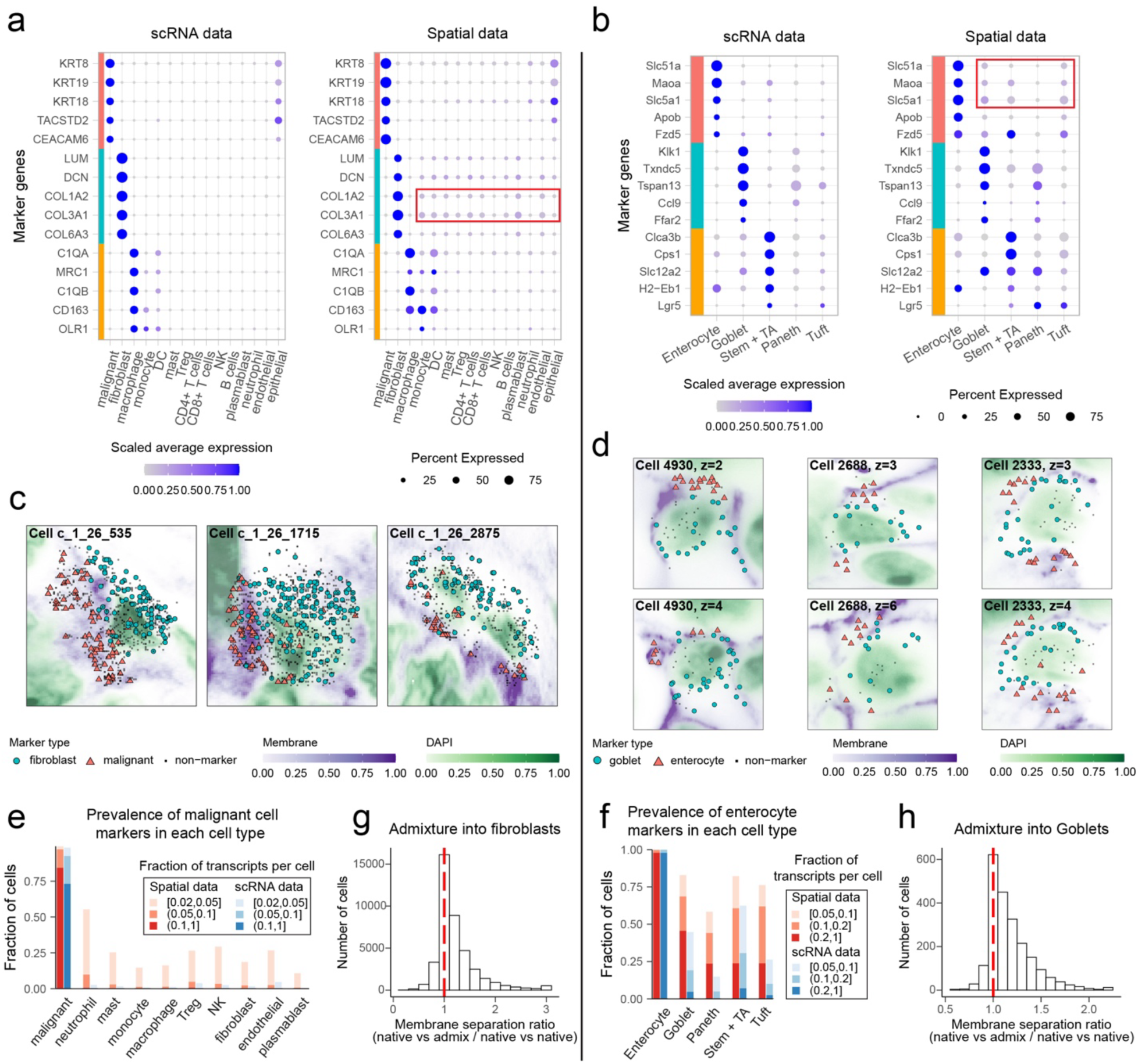
Segmentation errors cause admixture of molecules between adjacent cells. a. NSCLC cell type markers. Dot plot of top scRNA-seq cell type marker relative gene expression in scRNA-seq data (left) and the corresponding spatial dataset (right). b. As with (a) but for the mouse ileum dataset and corresponding scRNA-seq. c. Three segmented fibroblast cells, showing all molecules assigned to each cell. Color and shape indicate if the molecule is a marker gene identified in the scRNA-seq data (corresponding to the dot plot in a). Green background corresponds to DAPI stain intensity, and purple background corresponds to membrane stain intensity. d. As with (e) but for three goblet cells from the mouse ileum dataset. Two separate z-slices are shown for each cell. e. Comparing the dataset-wide prevalence of malignant cell admixture quantity into each cell type for the spatial data (red) versus scRNA-seq data (blue). Admixture quantity (color intensity) is calculated as the percent of a total cell’s transcripts coming from malignant marker genes (identified in scRNA-seq). The fraction of cells of a given type with each admixture degree is shown on the y-axis. f. As with (c) but for admixture of enterocyte markers into other cell types in the mouse ileum. g. Distribution of membrane separation ratio scores for fibroblasts in the NSCLC dataset. This ratio is greater than 1 if there is more membrane signal between native and admixture molecules compared to that between sets of only native molecules. h. As with (g) but for goblet cells in the mouse ileum.

Examining examples of cells with noticeable expression of foreign markers, we found that these potential admixture molecules often occurred in clusters close to the periphery of the cell (Figure 1c,d). In some cases, clusters of foreign markers were only found close to the top or bottom z planes in a given cell (Figure 1d). This indicates that admixture can come from cells in adjacent z-planes as well as in the same z-plane, highlighting the importance of considering segmentation accuracy in three dimensions. The general frequency and magnitude of such potential admixtures varied between datasets and cell types, but were higher than that observed in the scRNA-seq data (Figure 1e,f). This trend was also observed in the Xenium pancreas dataset when compared to a pancreas scRNA-seq dataset (Supp. Figure 1c).

We reasoned that the occurrence of such foreign markers at the cell periphery likely reflects mis-assignment of molecules during cell segmentation. To confirm this, we tested whether these potential admixture molecules tend to be separated from the molecules expected of the native cell type by a membrane signal. Modern cell segmentation methods use membrane stains to establish likely cell boundaries, so cases where the membrane signal was very clear were likely already segmented correctly. Nevertheless, if the expression of foreign markers represents segmentation errors, we expect to see some residual membrane signal separating these admixture clusters from the native transcripts. Indeed, we find that in both datasets where membrane stain was available, the average membrane signal along the paths connecting foreign and native marker gene molecules was significantly higher than it was along the paths connecting native markers (Figure 1g,h). Together these results indicate that mis-assignment of molecules from distinct cell types is prevalent in the current spatial datasets.

### Admixture artifacts dominate context-dependent cell expression differences

To evaluate the impact of such molecular admixtures on downstream analysis we first examined the simplest context-dependency test: differential expression of a given cell type between distinct tissue contexts. Such a test, for example, could be used in cancer studies to understand what transcriptional features distinguish immune cells infiltrating the tumor relative to cells positioned on the tumor boundary or within the adjacent normal regions.

As a first example, we compared the transcriptional state of the fibroblasts positioned within the NSCLC tumor regions with the fibroblasts found in the stromal regions (Figure 2a). The cell type composition of the two regions was markedly different: the tumor regions were dominated by the malignant cells, while the stromal regions were composed mostly of immune, fibroblast, endothelial and epithelial cell populations (Figure 2b). When carrying out a differential expression (DE) test on the fibroblast populations in the two regions, we found that the list of genes reported as upregulated in the tumor context was dominated by the malignant expression markers, including *KRT19* and *KRT8* among others. (Figure 2c). According to the scRNA-seq data, these classical epithelial markers should not be expressed in fibroblasts (Figure 1a). Overall, such malignant cell specific markers constituted 11 out of 36 top upregulated DE genes, with a GSEA enrichment p-value of 2.8x10^-4^. In contrast, genes reported as upregulated in fibroblasts positioned within the stromal regions included markers of other non-malignant populations, such as macrophages (*C1QA*, *CD163*).

**Figure 2.**
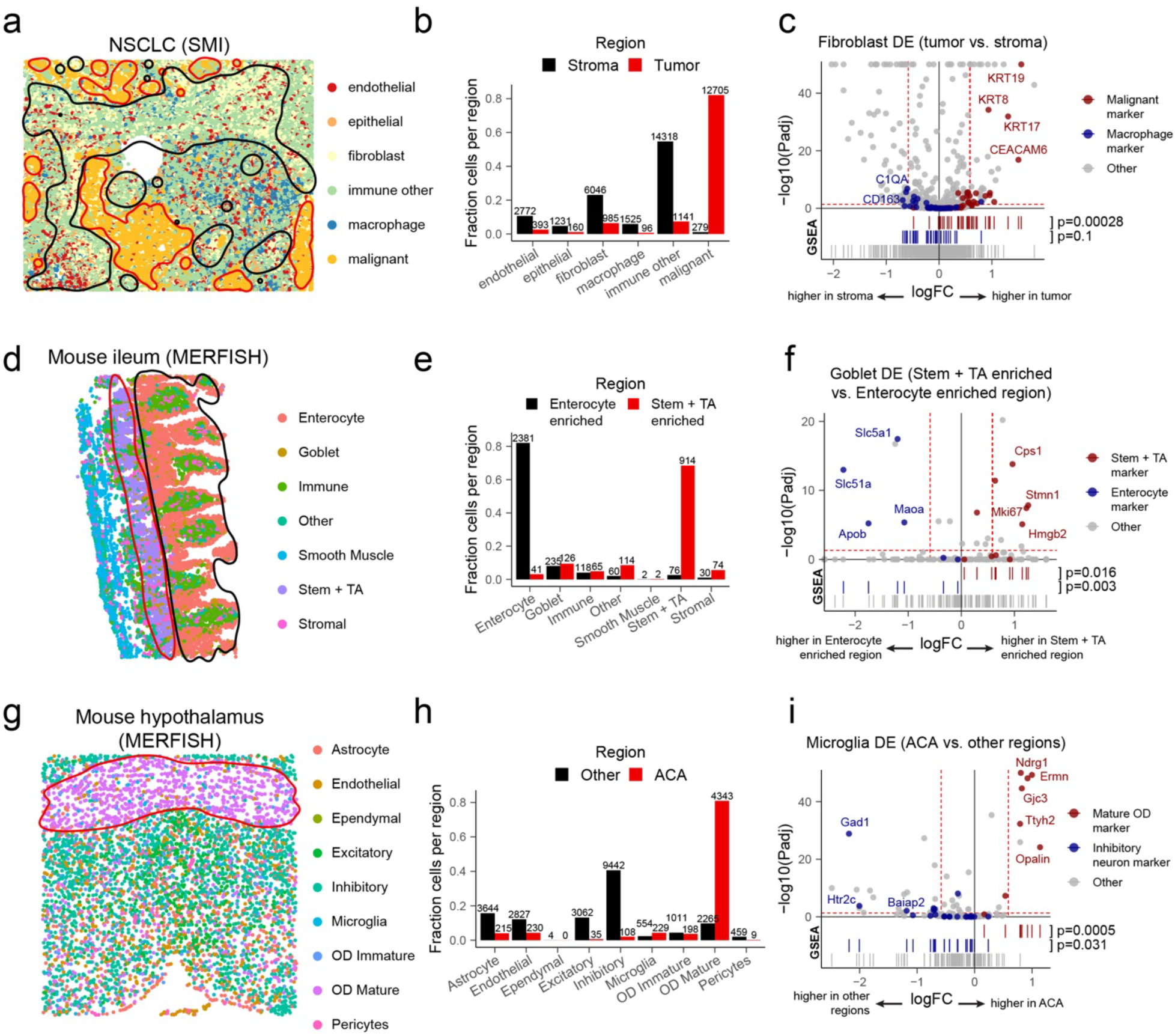
Segmentation errors confound context-dependent differential expression results. a. CosMx data from one NSCLC patient. Cell types are shown as different colored points. Circled regions indicate which contexts were used for comparison in differential expression. The tumor interior region is circled in red and the stroma is circled in black. b. Cell type composition in the tumor interior and stroma regions. Composition is shown as the fraction of all cells in a region coming from cells of a given cell type. c. Differential expression of fibroblasts between tumor and stroma regions. scRNA-seq marker genes of malignant cells and macrophages are shown as red and blue colors, respectively. The p-value is from GSEA to test for enrichment of marker genes among top DE genes. d. As with (a) but for the mouse ileum data. The stem + TA enriched crypt region is circled in red, and enterocyte enriched villi region is circled in black. e. As with (b) but for the mouse ileum crypt and villi regions. f. As with (c) but for goblet cell differential expression between crypt and villi regions. scRNA-seq marker genes of stem + TA cells and enterocytes are shown as red and blue colors, respectively. g. As with (a) but for the mouse hypothalamus dataset. The ACA region is circled in red. h. As with (b) but for the mouse ileum dataset ACA regions and all other regions. i. As with (c) but for microglia differential expression between the ACA and other regions. scRNA-seq marker genes of mature oligodendrocytes and inhibitory neurons are shown as red and blue colors, respectively.

Next, we observed analogous effect with the mouse ileum dataset, comparing expression in Goblet cells between crypt and villus regions (Figure 2d). Crypts are enriched with stem cells and transit-amplifying (TA) cells, while the villi are enriched for enterocytes (Figure 2e). Goblet cells found in the crypt regions had apparent upregulation of stem cell and TA markers (GSEA p-value = 0.016), while those in the villi had apparent upregulation of enterocyte markers (GSEA p-value = 0.003) (Figure 2f). In a third example, we observed this problem in the mouse hypothalamus dataset. Here, we compared expression of microglia cells in the anterior commissure (ACA) region versus those outside of this region. The ACA is enriched with mature oligodendrocyte (OD) cells (Figure 2h), and we observed an increased in mature OD marker expression in the microglia in this region (GSEA p-value = 0.0005) (Figure 2i).

Overall, these results show that molecular admixtures, likely stemming from segmentation errors, often dominate the results of the differential expression analysis between regions, obscuring true differences of cellular state.

### Admixture artifacts are also dominant in analysis of cell interactions and coordination

Different approaches have been developed to infer potential interactions or coordination between different cell types within a tissue^26–35^. For example, one may infer that two cell types interact if the state of a cell is consistently affected by the nearby presence of the other cell type^27,34^. The resulting expression changes are often termed “interaction changed genes”. To illustrate the impact of the admixture on such inference, we used a simple differential expression test to compare the state of macrophages positioned next to fibroblasts with those that do not have a fibroblast next to them (Figure 3a). Since segmentation errors tend to misassign molecules between adjacent cells, we hypothesized that we would observe an increase in apparent fibroblast marker gene expression for the group of macrophages located adjacent to fibroblasts compared to all others. Indeed, we found that the resulting list of DE genes is dominated by fibroblast markers, such as *COL1A2* and *COL3A1* (GSEA p-value = 0.00029) (Figure 3b). Similarly, we found an enrichment of contaminating marker genes in the interaction-changed genes from the pancreas dataset when testing endothelial cells near versus not near stellate cells (Supp. Figure 1d).

**Figure 3.**
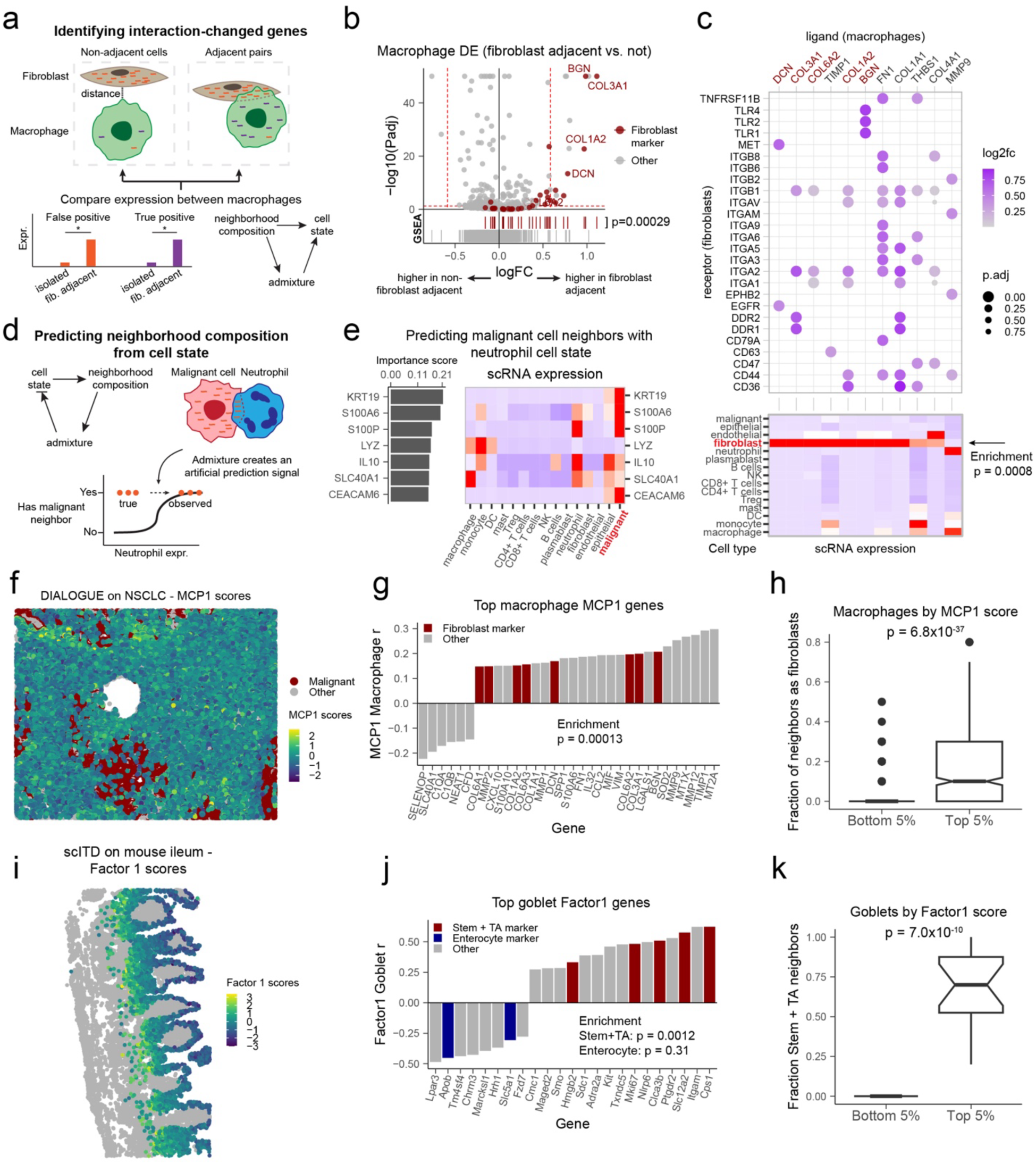
Segmentation errors can cause false positive ligand-receptor inferences and confound multi-cell type expression covariation across spatial regions. a. Diagram illustrating how interaction-changed genes are identified and how segmentation errors can lead to false-positives. b. Interaction-changed genes for macrophages. DE was run for macrophages which are adjacent to fibroblasts (N = 6048) versus those that are not adjacent to fibroblasts (N = 15,206). scRNA-seq marker genes from fibroblasts are shown in red. The p-value is from GSEA to test for enrichment of fibroblast marker genes among the top DE genes identified in macrophages. c. Macrophage to fibroblast ligand-receptor interaction results (top). Ligand genes called as fibroblast markers from scRNA-seq data are colored in dark red. Relative scRNA-seq gene expression of the ligand genes is shown in each cell type (bottom). d. Diagram illustrating a basic approach to predicting the cell type neighborhood composition from a given cell’s gene expression state. e. Results from predicting presence of a malignant cell neighbor from neutrophil expression state. Importance scores indicate which genes in neutrophils were the strongest predictors of malignant cell presence (left). Relative scRNA-seq gene expression for the top genes is shown across cell types (right). f. DIALOGUE MCP1 scores shown for one NSCLC patient. These scores were computed using fibroblasts, macrophages, and CD4+ T cells, and thus the scores for only those cell types are shown. Malignant cells are colored in red. g. Macrophage gene expression-MCP1 correlations for the DIALOGUE results on NSCLC data. scRNA-seq marker genes of fibroblasts are shown in red. The enrichment p-value is from GSEA test. h. The fraction of macrophage neighbors that are fibroblasts, shown separately for the macrophages with the bottom and top 5^th^ percentile MCP1 scores (N = 2718 total). The p-value is from a two-sample Student’s t test (two-sided). The notches on the boxplot indicate a 95% confidence interval around the median. i. scITD Factor1 scores for the mouse ileum data. These scores were computed using goblet cells and enterocyte cells, and thus the scores for only those cell types are shown. j. Goblet cell gene expression-Factor1 correlations for the scITD result on the mouse ileum data. scRNA-seq marker genes of Stem + TA cells are shown in red and marker genes of enterocytes are in blue. The p-value is from GSEA test. k. The fraction of goblet cell neighbors that are stem + TA cells, shown separately for the goblet cells with the bottom and top 5^th^ percentile Factor1 scores (N = 298 total). The p-value is from a two-sample Student’s t test (two-sided). The notches on the boxplot indicate a 95% confidence interval around the median.

Another approach to evaluate the impact of the cellular microenvironment is to construct a predictive model that evaluates the relationship between cellular state and the composition of the cell’s surrounding neighborhood^34,35^. To illustrate this, we trained a simple logistic regression model to predict whether neutrophils had a malignant neighbor based on their transcriptomic profile. This neighbor prediction model performed significantly better than chance on unseen test data (empirical p-value=10^-5^), suggesting that the presence of a malignant neighbor has a substantial transcriptional influence on neutrophils. However, when we inspected the top informative genes for the model we found that most of them (4 out of 7, *e.g. KRT19*, *CEACAM6*) were strongly expressed in malignant cells but not in neutrophils (Figure 3c). This is consistent with the hypothesis that segmentation errors, which misattribute some molecules from malignant cells to adjacent neutrophils, generate artifactual signal in predicting association between cell state and adjacent presence of another cell type. A similar pattern was observed when predicting fibroblast presence based on macrophage expression, where fibroblast markers were among the most important genes (Supp. Figure 2f).

A variety of computational methods have been developed to prioritize potential channels of inter-cellular communication through analysis of cognate ligand-receptor (LR) pairs^27,28,31–33,35^. In analysis of spatial data one could, for example, compare expression of cognate ligand and receptor genes in pairs of heterologous cell types when those cell types are adjacent to each other versus when they are apart. Since segmentation errors are more likely to misassign molecules between adjacent cells, such analysis approaches may result in false positive LR inferences if any ligand or receptor genes are highly expressed in one or both cell types (Figure 3d). To illustrate this, we used the Giotto LR inference method to identify potential LR signals from macrophages (ligand cell type) to fibroblasts (receptor cell type). We observed that the top predicted LR interactions were mostly driven by ligand genes, which, despite being attributed to macrophages, were specific fibroblast markers, including well-known *COL6A3*, *BGN*, or *DCN* genes (Figure 3c). Such false positive inferences occurred more frequently among the top LR results than expected by chance (hypergeometric test p-value = 7.7x10^-5^).

A distinct set of methods has been developed to identify patterns of transcriptional co-variation of multiple cell types across spatial niches. Two such tools include DIALOGUE^30^ (Jerby-Arnon 2022) and scITD^36^. Each method produces a set of factors (referred to as “multicellular programs” (MCPs) for DIALOGUE) that describe which genes in each cell type are correlated together across niches. Such transcriptional patterns may reflect niche-specific extracellular signals as well as intercellular interactions that vary across niches. In the presence of segmentation-errors, however, we may expect admixture signals to influence the resulting gene patterns. This is particularly likely when cell type composition changes across niches, as admixture signals will likely represent a dominant axis of variation in such cases. We analyzed the results of DIALOGUE applied to the NSCLC dataset (Figure 3f) and identified a significant enrichment of fibroblast markers among the top MCP1 genes for macrophages (p-value = 0.00013) (Figure 3g). Overall, such false positive inferences appeared to constitute 25% of the top 32 results. We further showed that the macrophages with high MCP1 scores were significantly more frequently neighboring fibroblasts (p-value = 6.8x10^-37^), supporting the hypothesis that cell type admixture has confounded the MCP in this case (Figure 3h). We also found a similar effect when using our own scITD tool, as applied to the mouse ileum dataset. Here, the top factor-associated genes in goblet cells were actually Stem and TA marker genes (enrichment p-value = 0.0012) (Figure 3j). In this case, false positive inferences appeared to constitute 30% of the top 23 results. We also found that the goblet cells with lower scores had significantly more frequent adjacencies to Stem and TA cells (p-value = 7.0x10^-10^), pointing to admixture as a confounder for this factor.

### Factorization of molecular neighborhoods enables the separation and correction of admixture artifacts

Since the admixture transcripts have composition distinct from the expression profile of the native cell type and tend to form spatial clusters within cells (Figure 1a,e), we reasoned that it should be possible to identify and correct for the admixture patterns by analyzing local molecular neighborhoods. We therefore sought to extend the analysis of molecular neighborhood composition vectors (NCVs)^22^ to subcellular levels. Examining NCVs of all molecules within a specific cell type, we applied weighted non-negative matrix factorization (NMF) to identify recurring subcellular patterns (Figure 4a; Methods). We annotated the resulting components based on the expression of different cell type markers (Figure 4b; Supp. Figure 2a). In the NSCLC dataset fibroblast cells, this approach uncovered a factor (factor 2) driven by tumor markers, reflecting misassigned molecules from adjacent malignant cells (Figure 4b). Another factor (factor 1) was driven by macrophage markers (Supp. Figure 2a), reflecting potential admixture of molecules from adjacent immune cells into fibroblasts. The analysis also identified factors that separated the cytoplasm (factor 5) and nucleus (factor 4) compartment composition of the native fibroblast transcripts (Supp. Figure 2a). For other cell types where we observed apparent admixture, the weighted NMF approach again successfully identified factors which clearly correspond to markers of other cell types (Supp. Figures 3a, 4a, 5b, and 1*e).

**Figure 4.**
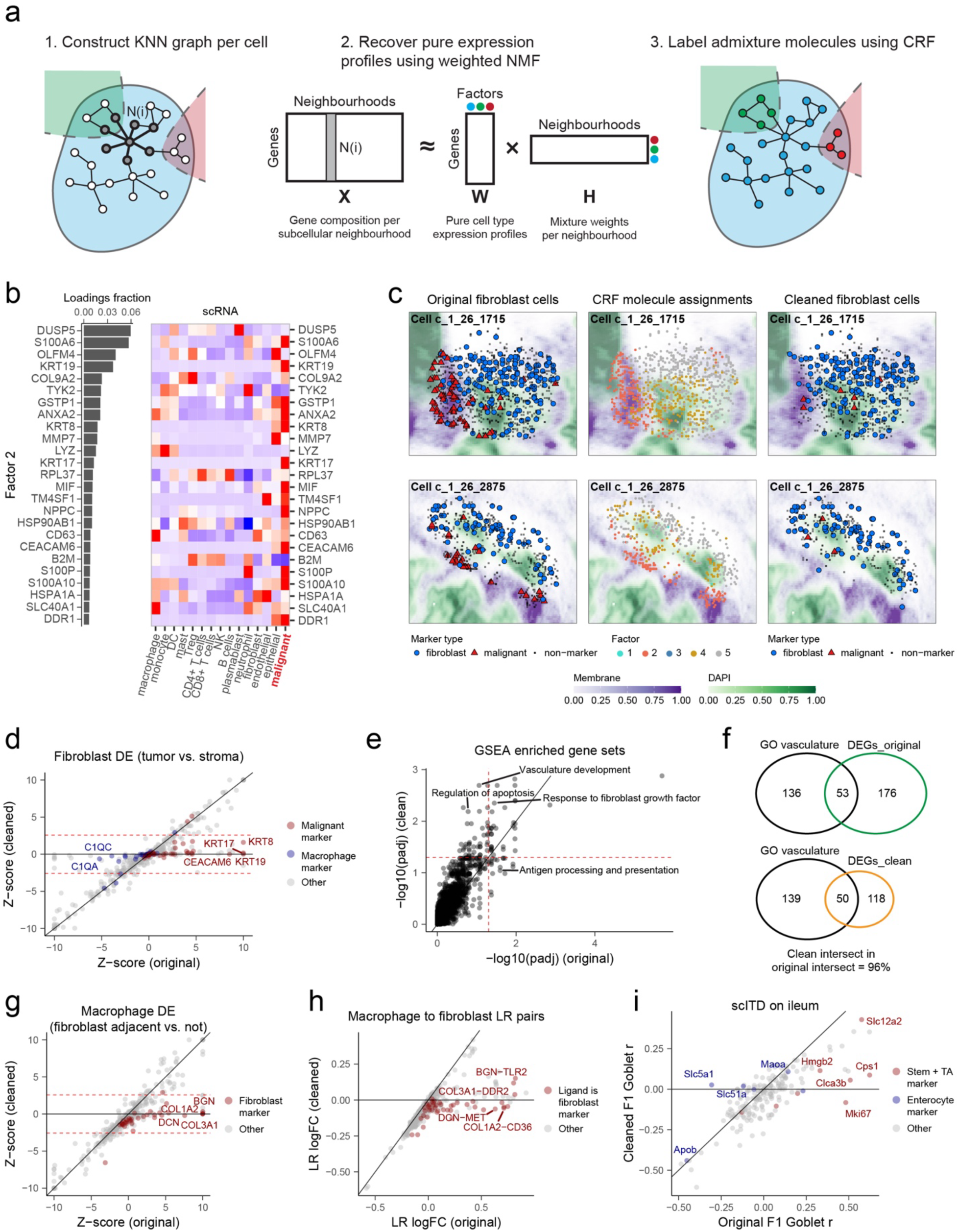
Factorization-based correction method minimized admixture from segmentation-errors while preserving biological variability. a. Diagram of the factorization of molecular neighborhoods used for corrections. b. The factorization applied to fibroblasts from the NSCLC data. (left) Gene loadings for Factor 2, capturing malignant cell admixture are shown. (right) Relative scRNA-seq gene expression of these top genes across cell types. c. Two segmented fibroblast cells, showing all molecules assigned to each cell. In the left plots, color and shape indicate if the molecule is a marker gene identified in the scRNA-seq data. In the middle plots, molecules are colored by their CRF factor assignments. The rightmost plots show the cells after removing molecules assigned to the malignant factor, Factor 2. d. Impact of data cleaning on fibroblast DE for tumor versus stroma regions. scRNA-seq marker genes of malignant cells are shown in red. e. Impact of data cleaning on the GSEA of the DE results shown in (d). f. Overlap of the GO_vasculature gene set with significant fibroblast DE genes (Padj < 0.01) either pre or post cleaning. g. As with (d) but for macrophage DE in the scenario of identifying fibroblast interaction-changed genes. scRNA-seq marker genes of fibroblast cells are shown in red. h. Impact of cleaning on macrophage to fibroblast LR pairs. Showing the LR log fold change from Giotto, which reflects the relative LR pair expression score for proximal versus distal cell pairs. A given LR pair is colored red if the ligand was a marker gene of fibroblasts identified from the scRNA-seq data. i. Impact of cleaning on scITD results. Showing goblet expression-Factor1 associations for pre versus post cleaning. scRNA-seq markers of stem + TA cells are colored red and markers of enterocytes are colored blue.

Given the spatially-clustered nature of the admixing patterns, we used a Conditional Random Field (CRF) model to assign individual molecules to the components uncovered by NMF. Examples of such molecular assignment are shown in Figure 4c, where in fibroblasts the admixture factor (factor 2) correctly selects the cluster of genes enriched for malignant marker genes. We found that the distribution of factor assignments was highly variable across spatial regions (Supp. Figure 2c), reflecting the differential abundance of the contamination source cell type across tissue compartments. For example, when inspecting the distribution of molecules assigned to the fibroblast factor 2 (malignant admixture factor) we found, as expected, that it had a larger fraction of molecules in tumor regions compared to those assigned to other factors (Supp. Figure 2c).

Notably, the removal of molecules assigned to admixture components minimized the impact of admixtures on the differential gene expression analysis, as reflected by the reduced number of foreign marker genes that appeared as significantly differentially expressed (Figure 4d, g). For fibroblast differential expression between tumor and stroma regions (NSCLC data), for example, we originally identified many malignant marker genes upregulated in fibroblasts in the tumor region. After applying our NMF and CRF clean-up procedure, we find that malignant markers are no longer statistically significant in the fibroblast DE analysis (Figure 4d). This indicates that our procedure successfully removed such admixture molecules from fibroblasts in the tumor region, which was confirmed by inspecting expression levels of these markers per region (Supp. Figure 2d). Importantly, while our approach removed the suspected admixture transcripts, it did not significantly impact expression of the native fibroblast marker genes (Supp. Figure 2e).

The removal of admixture molecules also had the effect of removing likely false positive enriched gene sets from GSEA run on the DE results. For example, the originally significant “antigen processing and presentation” gene set in fibroblasts was likely due to increased admixture of macrophages into fibroblasts outside of the tumor region (Figure 2c) and was no longer significant after applying our correction (Figure 4e). Additionally, we also obtained several newly significant gene sets after removing the admixture transcripts, which mostly involved vasculature development genes in the fibroblasts (Figure 4e and 4f). In general, these newly significant gene sets were not reported as significant in the original data (before clean-up) because the presence of many admixture molecules increased the number of total genes tested without adding more genes to the intersection with the gene sets, effectively diluting the signal (Figure 4f). These results suggest that fibroblasts in the tumor region may play a different role in angiogenesis compared to fibroblasts in the stroma. Overall, the removal of contaminating transcripts helped to reveal an interesting and relevant biological signal that would have otherwise been obscured. Our correction procedure also successfully alleviated admixture signals appearing in the other DE analyses of goblet cells in the mouse ileum dataset (Supp. Figure 4d), microglia in the mouse hypothalamus dataset (Supp. Figure 5e), macrophage-fibroblast adjacency DE in the NSCLC dataset (Figure 4g), and endothelial-stellate adjacency DE in the pancreatic cancer dataset (Supp. Figure 1g).

Removal of contaminating transcripts also diminished the number of such markers appearing in the top LR results and scITD factors (Figure 4h and 4i). For example, we had originally observed a significant enrichment of fibroblast genes in macrophage ligands of significant LR pairs for macrophage-to-fibroblast interactions. After applying the correction, the fold-change of LR interaction scores for these likely false LR pairs is greatly reduced (Figure 4h). Only two ligands, both of which had notable expression in macrophages, remained significant after correction (Supp. Figure 3e). These were *THBS1* and *MMP9*, each interacting with several possible receptors in the fibroblast cells. In correcting expression in goblet cells (mouse ileum dataset), the factorization procedure similarly mitigated the effects of admixture on scITD results. Originally, the top loading genes in goblet cells were enriched for Stem + TA marker genes. After correction, however, the scITD factor associations with these genes was greatly reduced, indicating they are less influential on the factor (Figure 4i). The correlation between the original and cleaned scITD factor scores was 0.93, indicating that the overall expression variation captured by the factor remained largely unchanged despite reducing the influence of these admixture genes.

## Discussion

Our study demonstrates that molecular admixtures resulting from imperfect cell segmentations are prevalent in imaging-based spatial transcriptomics data. In many of the downstream analyses these artifactual signals dominated the results, often constituting the majority of the top genes. This was true irrespective of the segmentation method used, and was apparent even in datasets with relatively clean segmentations, such as NSCLC or the Multimodal Cell Segmentation of the pancreas data. The overall impact of such molecular misassignments is akin, though likely weaker, to the convolution of transcriptional signals seen in high-resolution sequencing-based methods such as Slide-seq2 or Stereo-seq. While imaging-based data offers more tractable means for addressing such admixtures, most current analysis methods do not account for these errors.

The proposed correction method relies on factorization of neighborhood composition vectors (NCVs). We have previously demonstrated that at lower resolution, NCVs are effective at capturing transcriptional composition of distinct cell types and can be used for segmentation-free analysis^22^. In this study we showed that higher resolution NCVs, when combined with a robust factorization method, can distinguish recurrent subcellular features based on their mRNA composition. These features included true cellular structures, such as nuclei, endoplasmic reticulum, or cell polarization (Supp. Figure 6). However, most factors appeared to capture recurrent admixture patterns - mis-segmented patches of molecules from other cell types.

While we show that controlling for the admixture components can largely mitigate the impact of segmentation errors in common downstream analysis tasks, further developments are merited. Several aspects of the current procedure could be automated, most notably the classification of admixture factors. In our analysis we manually classified admixture factors based on the prevalence of foreign cell type markers determined from reference scRNA-seq data. This process can be automated and even integrated with large reference atlases. However, the reliance on scRNA-seq profiles is complicated by the presence of potential doublets in that data, as well as the fact that the admixture composition may not always correspond to whole-cell transcription profiles. Instead, it may reflect distinct cellular compartments such as cytoplasm, leading to systematic deviations from the scRNA-seq. Other characteristics, such as spatial coherence or similarity to neighboring cells, may also aid in recognizing admixture factors. The accuracy of the correction procedure may also be improved by performing factorization on all cell types simultaneously, though improvements in factorization stability and automated selection of the number of factors will be necessary.

Some elements of our correction procedure can undoubtedly be incorporated into cell segmentation algorithms themselves. However, attaining perfect segmentation will likely remain elusive. Similarly, we expect some residual admixture signal to remain even after applying the presented correction procedure. We think, therefore, that it will be important to extend downstream analysis methods to explicitly control for segmentation errors^9^. Overall, we hope that raising awareness of the confounding effects of segmentation errors, along with the suggested correction method, will enable to uncover molecular mechanisms orchestrating tissue biology.

## Methods

### Dataset processing

First, we needed to ensure that our spatial and scRNA-seq datasets contained the same cell types and were annotated in the same manner. Therefore, we plotted each spatial dataset and its corresponding scRNA-seq dataset together in a co-embedding using Conos^37^. We kept all clusters which visually overlapped and renamed each cluster to be consistent across datasets. For the NSCLC CosMx data, we only used donor 5 replicate 1 for membrane separation analysis, admixture prevalence analysis, and NMF. For all other downstream analyses including CRF, we used all three replicates for donor 5. For the mouse ileum MERFISH dataset, we selected only molecules with a confidence score greater than 0.5. For the mouse hypothalamus MERFISH dataset, we used animal ID = 1 for NMF and all female, naive-behavior mice for downstream analyses. We used the subset tissue slices corresponding to Bregma values of 0.16 and 0.21, as these contained the region of interest for DE analysis.

### Context-dependent differential expression

In order to run differential expression across discrete spatial contexts, we either relied on previously annotated contexts or computed them ourselves. For the NSCLC dataset, we utilized the existing niche annotations, specifically selecting the regions labeled “tumor interior” and “stroma” for comparison. For the mouse ileum and mouse hypothalamus datasets, we computed spatial niches by clustering the cell type k-nearest neighbor (KNN) graph, as in He et al. 2022^25^. We used KNN parameters of 10 and 8 for the ileum dataset and hypothalamus datasets, respectively. For the hypothalamus dataset, we ran KNN separately for each mouse and tissue slice but clustered all KNN vectors together. We only used the tissue sections at Bregma units of 0.16 and 0.21 as these contained the largest fraction of cells belonging to the ACA region, which was our selected region of interest. For the ileum dataset we selected the clustered regions belonging to the intestinal crypts and outer villi for comparison. For the hypothalamus region, we selected the clustered regions belonging to the ACA and all other regions for comparison. Next, differential expression was run across the selected regions for a given cell type by using the R package Pagoda2 (https://github.com/kharchenkolab/pagoda2). We report the log2 fold change values and p-values adjusted with the Benjamini-Hochberg procedure. GSEA p-values for enrichment of scRNA-markers among DE genes were computed with the *fgsea* R package, using the DE fold changes as the input metric. To test for enriched gene ontology (GO) gene sets, we applied *fgsea*, using the absolute value DE Z-scores to mitigate the impact of admixture signal on the results.

### Intercellular interactions

To test for interaction-changed genes in the NSCLC dataset, we first stratified macrophages into two groups. One group consisted of macrophages that had at least one fibroblast cell within its 10 nearest neighbors. The second group consisted of macrophages without any fibroblast neighbors. Then, we ran DE and GSEA as above. To identify ligand-receptor interaction between macrophages and fibroblasts, we applied the method implemented in the Giotto R package^27^. We paired this with cognate LR pairs from the iTalk database^38^. We ran the inference with 1000 random permutations and adjusted for multiple hypothesis correction with FDR correction. To select the top results for visualization, we thresholded all results to ligands that were part of any interaction with an adjusted p-value < 0.05 and a log2 fold-change greater than 0.25. We further removed receptors from the heatmap if they had no significant interaction with any ligand. Among the top ligands identified in macrophages, we tested whether they were enriched for fibroblast marker genes (from scRNA-seq) using a hypergeometric test.

### Multicellular programs

We ran DIALOGUE for the NSCLC data using the same input data and parameters as in the original publication. This was done by following the tutorial in the DIALOGUE GitHub at: https://github.com/livnatje/DIALOGUE/wiki/Reproducing-the-lung-cancer-spatial-results-and-figures. To run our tool, scITD, on the mouse ileum dataset, we first created spatial patches, each containing at least one enterocyte and one goblet cell. This yielded 10 cells per patch on average. Expression per cell type per patch was pseudobulked and normalized using the scITD default parameters. Then, we ran scITD to extract three three factors (multicellular patterns of variation across patches).

### Marker gene selection

To select clean sets of cell type markers, we utilized scRNA-seq datasets of the same tissues matching the spatial panels. For NSCLC we used a scRNA-seq dataset from Zilionis et al. 2019 (accession GSE127465)^39^. For the mouse ileum, we used a dataset from Haber et al. 2017^40^. For the mouse hypothalamus, we used a scRNA-seq dataset which was paired with the spatial data in the original publication^24^. We used Seurat to identify markers from the NSCLC dataset and pagoda2 for the mouse ileum and hypothalamus datasets. For NSCLC, markers were chosen as having an adjusted p-value < 0.01, a log2FC > log2(1.5), and an AUC > 0.6. We further applied a filter to remove markers which were non-specifically expressed in multiple cell types. For the ileum data, markers were chosen as having an adjusted p-value < 0.05, and we again removed genes which were non-specifically expressed. For the hypothalamus dataset, markers were chosen as having an adjusted p-value <0.05 and a precision > 0.5. We further kept only genes that were found in each respective spatial marker panel.

### Neighbor prediction

Given a *focal cell type* 𝑐_𝑎_ and a *neighbor cell type* 𝑐_𝑏_, we are interested in assessing whether the presence of 𝑐_𝑏_ influences the transcriptome of 𝑐_𝑎_. We formulate this as a supervised learning problem.

Suppose we have 𝑁 cells of type 𝑐_𝑎_ with transcriptomes 𝑢_1_, 𝑢_2_, . . . , 𝑢_𝑁_. For 𝑘 ∈ {1, 2, . . . , 𝑁}, let 𝑦_𝑘_ ∈ {0, 1} denote whether or not cell 𝑘 has a cell of type 𝑐_𝑏_ in its neighborhood (which we define as its 6 nearest neighbors). We divide the data into training, validation, and test splits (see below) and fit a logistic regression model to approximate 𝑝(𝑦|𝑢). If the model performs better than chance on the test set, we conclude that the presence of 𝑐_𝑏_ is associated with transcriptional changes in 𝑐_𝑎_. Further, we can use the weights of the logistic regression model to rank the most important genes to the model.

We split the data into training (50%), validation (25%), and test (25%) splits. To control for spatial autocorrelation, we discretize the spatial domain into a 10x10 grid and split at the level of grid units (i.e. all cells in a grid unit are assigned to the same split).

The expression data is normalized before being passed to the model. First, the raw counts are normalized to sum to 1 for each cell. We then scale these values by 1e4, add a pseudocount, and take the natural log. We then compute the per-gene mean and variance on the training set and use those values to standardize the data in all three splits (i.e. subtract the mean and divide by the variance).

Each gene is assigned an “importance score” that indicates the extent to which it is a marker for the presence of a neighbor of cell type 𝑐_𝑏_. For simplicity, we use the value of the logistic regression parameter for each gene as its importance score. Each weight can be interpreted as the amount by which the log-odds ratio increases when the corresponding gene expression value increases by one unit.

All models are trained in PyTorch^41^. We use a binary cross-entropy loss. To manage the imbalance between positive and negative labels, instances with a positive label receive a weight equal to ∑(1 − 𝑦_𝑘_) / ∑ 𝑦_𝑘_, where this sum is computed over the training set. We use the Adam optimizer^42^ with a learning rate of 1e-4, a batch size of 64, and train for 300 epochs. Our final model is chosen to be the one that achieves the highest average precision on the validation set, which is checked after each training epoch.

To test whether a model performs better than chance, we use the average precision on the test set as our observed test statistic. Chance-level performance for average precision is equal to the fraction of instances with a positive label. We draw bootstrap samples to approximate the null distribution and compute an empirical p-value as the fraction of bootstrap samples whose statistic exceeds the observed test statistic.

### Membrane separation scoring

One way to validate admixture calls is to check whether they are consistent with membrane stains. Intuitively, molecules in a cell that are called as *admixture* should be separated from molecules called as *native* by an unusually high membrane stain intensity.

To make this idea concrete, let 𝑠(𝑚_1_, 𝑚_2_) denote the average membrane stain intensity along the straight line connecting two molecules 𝑚_1_ and 𝑚_2_. Define the pairwise maximum score

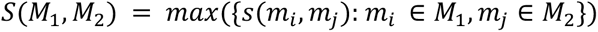

where 𝑀_1_ and 𝑀_2_ are disjoint sets of molecules. Suppose we are given a set of molecules 𝑀_𝑛𝑎𝑡𝑖𝑣𝑒,𝑘_ deemed to be native to cell 𝑘 and a set of molecules 𝑀_𝑎𝑑𝑚𝑖𝑥,𝑘_ deemed to be admixture of another cell into cell 𝑘. We compute the membrane separation ratio for cell 𝑘 as

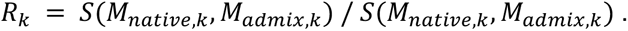

Normalizing by 𝑆(𝑀_𝑛𝑎𝑡𝑖𝑣𝑒,𝑘_, 𝑀_𝑎𝑑𝑚𝑖𝑥,𝑘_) is meant to account for the fact that membrane stains can be highly variable. If 𝑅_𝑘_ > 1 then the molecules called as admixture tend to be separated from native molecules by more membrane signal than separates native molecules from each other. We formally test whether 𝑅 = 𝑚𝑒𝑎𝑛({𝑅_𝑘_}_𝑘_) > 1 using a one-sided *t*-test.

Practically, we filter out any cell 𝑘 with 𝑚𝑖𝑛(|𝑀_𝑛𝑎𝑡𝑖𝑣𝑒,𝑘_|, |𝑀_𝑎𝑑𝑚𝑖𝑥,𝑘_|) < 5. Further, when computing 𝑆(𝑀_1_, 𝑀_2_) we only use pairs of molecules that lie in the same *z*-plane since there may be out-of-plane structures or membrane stain variability between *z*-planes. The sets 𝑀_𝑛𝑎𝑡𝑖𝑣𝑒,𝑘_ and 𝑀_𝑎𝑑𝑚𝑖𝑥,𝑘_ are derived from marker genes. In particular, if we have marker genes for each cell type of interest then we define 𝑀_𝑛𝑎𝑡𝑖𝑣𝑒,𝑘_ to be all molecules of marker genes for the cell type of cell 𝑘 and we define 𝑀_𝑎𝑑𝑚𝑖𝑥,𝑘_ to be all molecules corresponding to marker genes of any other cell type.

### Factorization and admixture effect correction

To detect intercellular admixtures from segmented spatial transcriptomic data, we first apply non-negative matrix factorization to recover subcellular expression patterns. Based on the existing cell segmentation, we construct subcellular neighborhood composition vectors (sNCVs) that reflect the abundance of transcripts from each gene near an RNA molecule. Different from the regular NCV as described in Petukhov et al.^22^, only neighboring molecules within cells are considered. In addition, the neighborhood size of sNCV is selected to be less than half of the average molecule count per cell in order to capture subcellular variations in transcript composition. Then, a weighted NMF algorithm (implemented in the R package “NMF”^43^) is applied to sNCVs of all RNA molecules from the same cell type^44^, where genes are inversely weighted by their average expression level. The resulting NMF components are examined for marker genes from other cell types. Components that are enriched with such marker genes are labeled as cross-cell-type contamination, while the rest are labeled as putative subcellular structures. Second, to assign individual transcripts to subcellular components while leveraging spatial information, we applied a Markov random field model on the KNN graph built from transcripts of individual cells. A transition matrix is defined such that the transition probability is α between compartments and 1-(t-1)*α within compartments, where α = 0.05 and t is the number of factors. The emission probabilities for gene labels of individual molecules are configured using a multinomial distribution, where the event probabilities correspond to the normalized (to sum to 1) gene weights in each NMF component. We used belief propagation implemented by R package “CRF” (https://github.com/wulingyun/CRF) to derive the posterior probabilities and maximum likelihood estimates of the compartment labels of individual molecules.

## Data availability

The NSCLC CosMx dataset is publicly available at https://nanostring.com/products/cosmx-spatial-molecular-imager/ffpe-dataset/nsclc-ffpe-dataset/. The NSCLC scRNA-seq dataset can be found at GEO accession GSE127465. The mouse hypothalamus MERFISH dataset is available at https://doi.org/10.5061/dryad.8t8s248. The mouse hypothalamus scRNA-seq dataset is available at GEO accession GSE113576. The mouse ileum MERFISH data can be found at https://doi.org/10.5061/dryad.jm63xsjb2. The mouse ileum scRNA-seq data can be found at GEO accession GSE92332. The pancreatic cancer Xenium data can be found at https://www.10xgenomics.com/datasets/ffpe-human-pancreas-with-xenium-multimodal-cell-segmentation-1-standard. The corresponding pancreas snRNA-seq dataset we used can be found at http://singlecell.charite.de/cellbrowser/pancreas/.

## Code availability

We provide supplementary files containing an R script with functions to run our CRF-based correction procedure as well as a tutorial notebook illustrating how to run it.

---

**Supplementary Figure 1.**
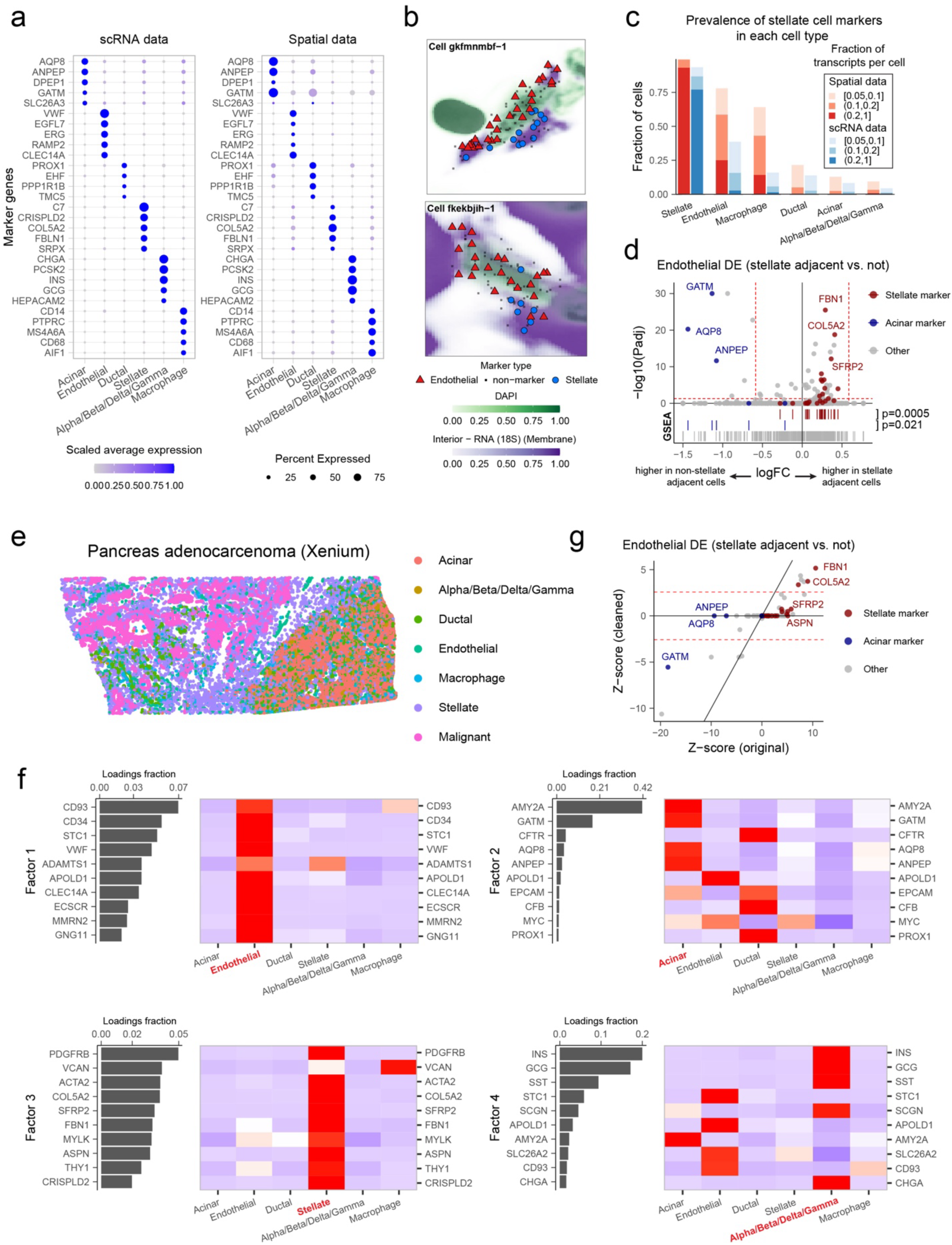
Segmentation error impact and correction for Xenium human pancreatic cancer dataset. a. Pancreas cell type markers. Dot plot of top scRNA-seq cell type marker relative gene expression in scRNA-seq data (left) or the corresponding spatial dataset (right). b. Two segmented endothelial cells, showing all molecules assigned to each cell. Color and shape indicate if the molecule is a marker gene identified in the scRNA-seq data for endothelial cells or stellate cells (source of admixture). Green background corresponds to DAPI stain intensity, and purple background corresponds to membrane stain intensity. c. Comparing the dataset-wide prevalence of stellate cell admixture quantity into each cell type for the spatial data versus scRNA-seq data. Admixture quantity (color) is calculated as the percent of a total cell’s transcripts coming from stellate cell marker genes (identified in scRNA-seq). The fraction of cells of a given type with each admixture degree is shown on the y-axis. d. Interaction changed genes for endothelial cells for those located near stellate cells (N = 862) versus those not near stellate cells (N = 608). scRNA-seq marker genes of stellate cells and acinar cells are shown as red and blue colors, respectively. The p-value is from GSEA to test for enrichment of marker genes among top DE genes. e. Xenium data from a pancreatic adenocarcinoma patient sample. Cell types are shown as different colored points. f. Top NMF factor loadings for NMF run on endothelial cells (left plots). Relative scRNA-seq expression is also shown (right plots). g. Impact of data cleaning on endothelial cell interaction changed genes DE. scRNA-seq marker genes of stellate cells and acinar cells are shown in red and blue colors, respectively.

**Supplementary Figure 2.**
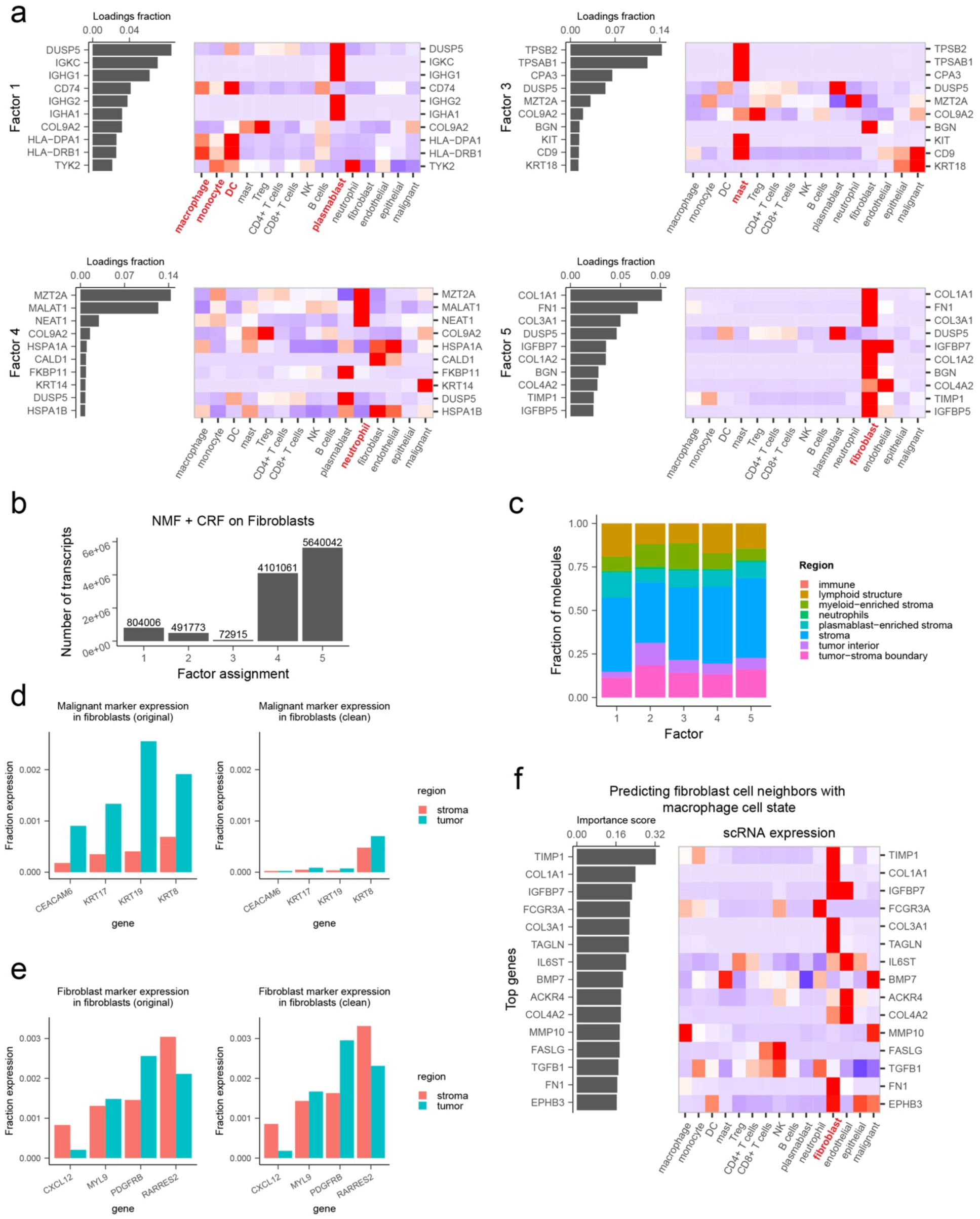
Additional NSCLC results for cleaning fibroblast cells. a. Top NMF factor loadings for NMF run on fibroblast cells (left plots). Relative scRNA-seq expression is also shown (right plots). Factor 2 is excluded here as it is shown in the main Figure 4b. b. Number of transcripts assigned to each factor. c. Fraction of all molecules assigned to a given factor that are found in each region. d. Expression of scRNA-seq selected malignant marker genes in spatial data fibroblasts (admixed genes) pre- (left) versus post- (right) cleaning. e. Expression of scRNA-seq selected fibroblast marker genes in spatial data fibroblasts (native genes) pre- (left) versus post- (right) cleaning. f. Results of predicting cell type neighborhood composition from cell state using macrophage gene expression to predict fibroblast neighbor presence. Importance scores indicate which genes in macrophages were strongest predictors of fibroblast cell presence (left). Relative scRNA-seq gene expression for the top genes is shown across cell types (right).

**Supplementary Figure 3.**
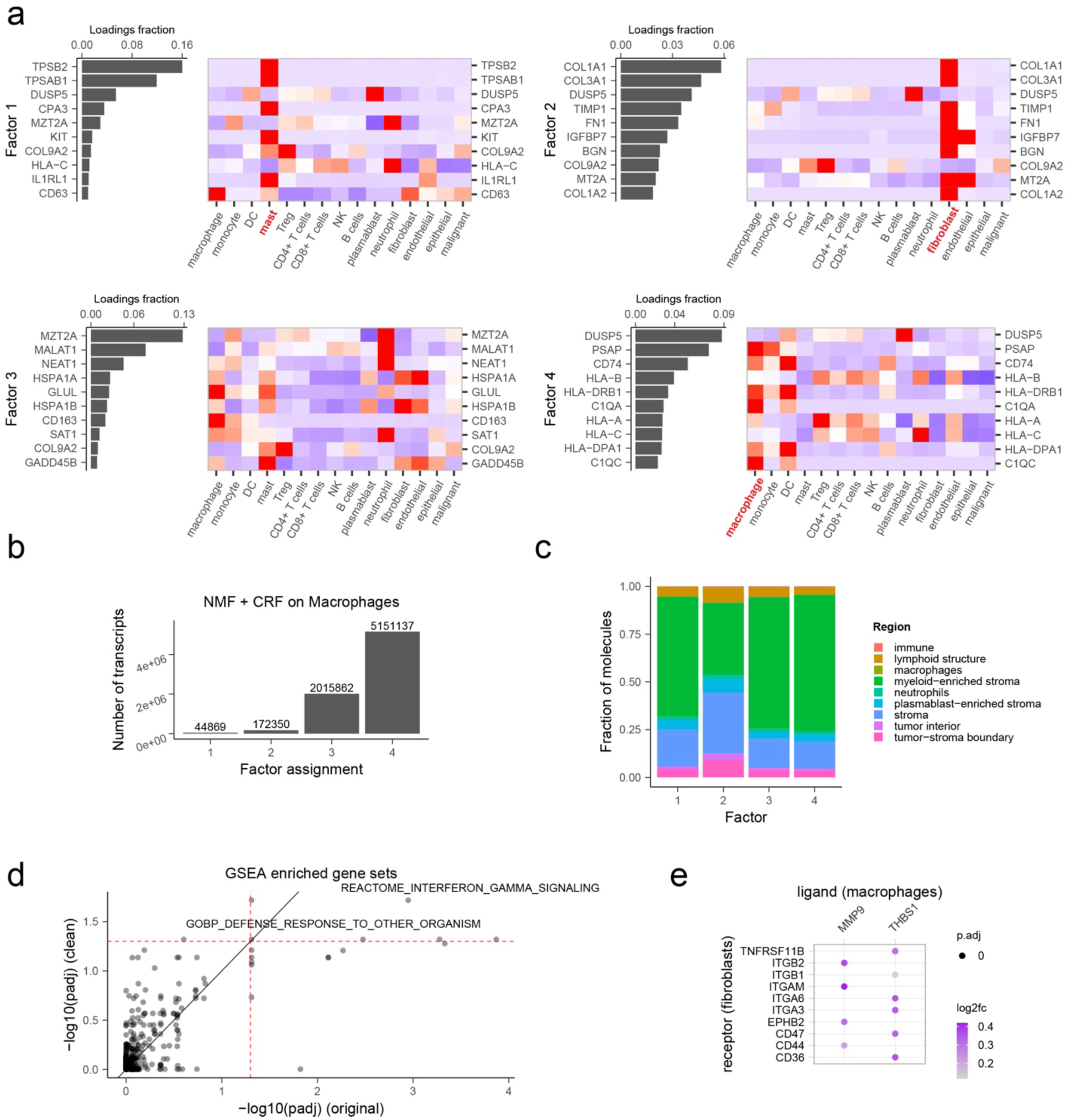
Additional NSCLC results for cleaning macrophage cells. a. Top NMF factor loadings for NMF run on macrophage cells (left plots). Relative scRNA-seq expression is also shown (right plots). b. Number of transcripts assigned to each factor. c. Fraction of all molecules assigned to a given factor that are found in each region. d. GSEA results for macrophage DE from fibroblast interaction changed gene pre versus post cleaning. e. Ligand-receptor inference results post cleaning. The same thresholds were used here as in Figure 3c.

**Supplementary Figure 4.**
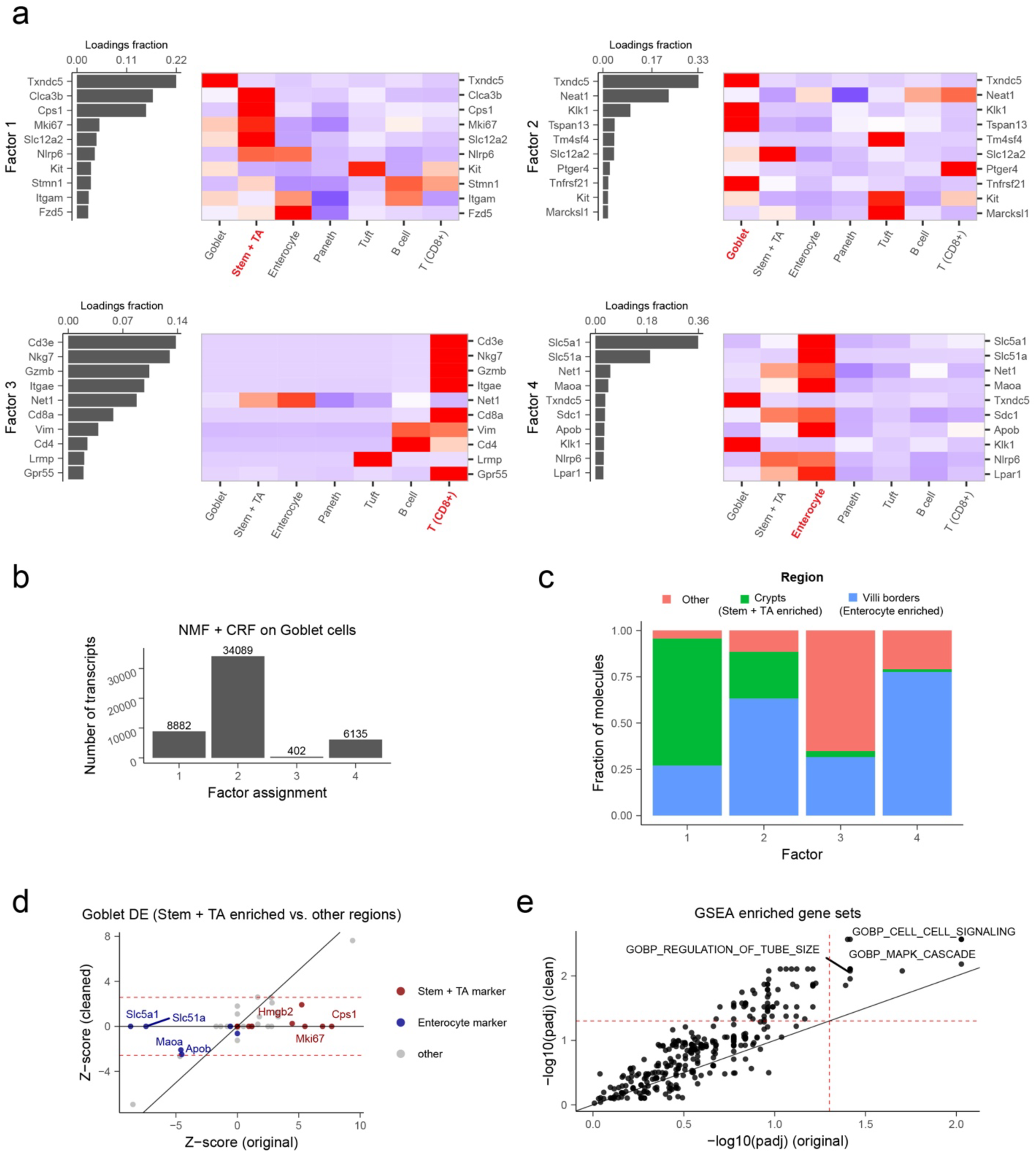
Additional mouse ileum results for cleaning goblet cells. a. Top NMF factor loadings for NMF run on goblet cells (left plots). Relative scRNA-seq expression is also shown (right plots). b. Number of transcripts assigned to each factor. c. Fraction of all molecules assigned to a given factor that are found in each region. d. Impact of data cleaning on goblet DE for crypt (Stem + TA enriched) versus villi (enterocyte enriched) regions. scRNA-seq markers of stem + TA cells are colored red and markers of enterocytes are colored blue. e. GSEA results for goblet DE between crypts versus villi.

**Supplementary Figure 5.**
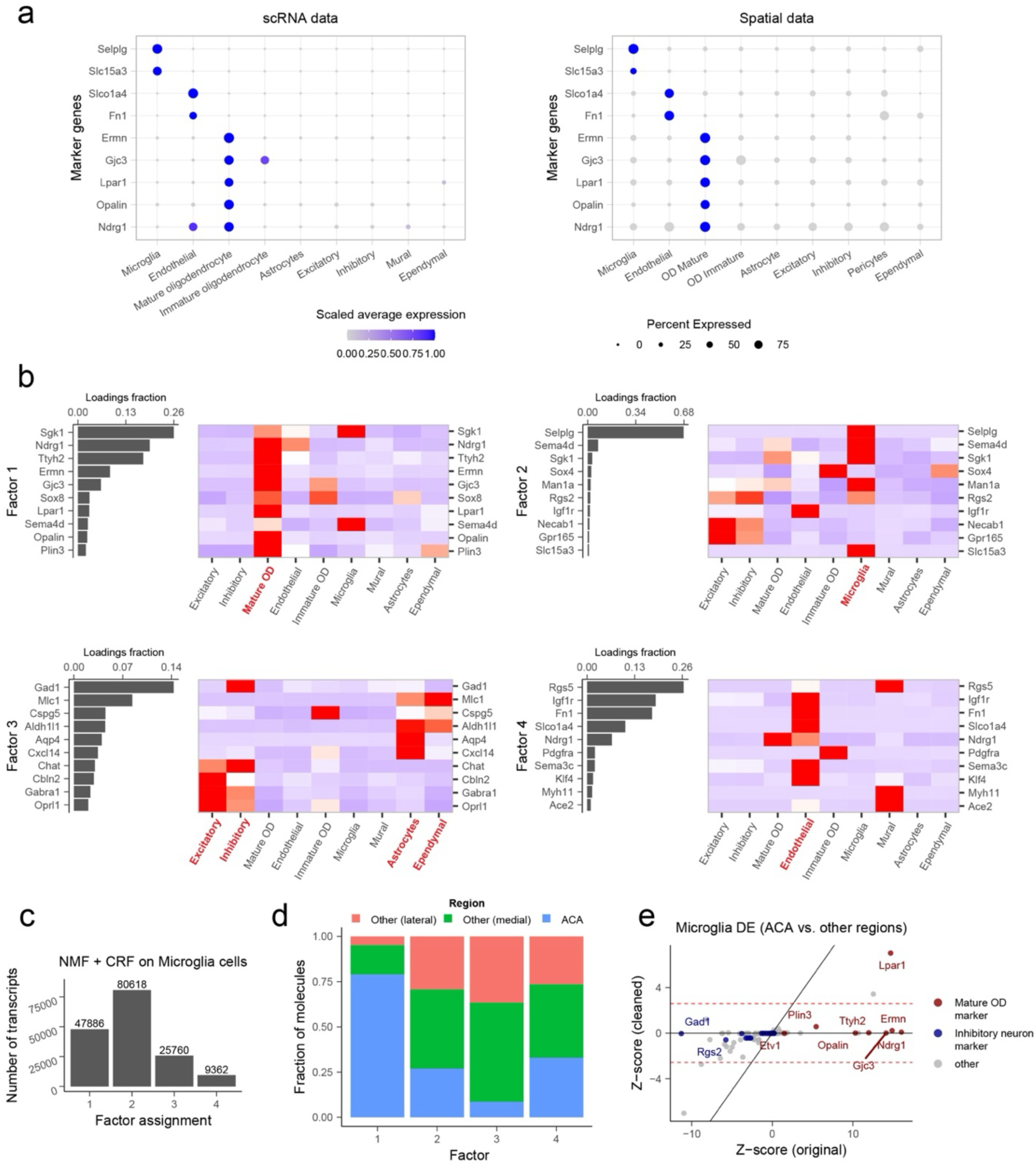
Additional mouse hypothalamus results for cleaning microglia cells. a. Mouse hypothalamus cell type markers. Dot plot of top scRNA-seq cell type marker relative gene expression in scRNA-seq data (left) or the corresponding spatial dataset (right). b. Top NMF factor loadings for NMF run on microglia cells (left plots). Relative scRNA-seq expression is also shown (right plots). c. Number of transcripts assigned to each factor. d. Fraction of all molecules assigned to a given factor that are found in each region. e. Impact of data cleaning on microglia DE for ACA versus other regions. scRNA-seq markers of mature oligodendrocytes and inhibitory neurons are colored red and blue, respectively.

**Supplementary Figure 6.**
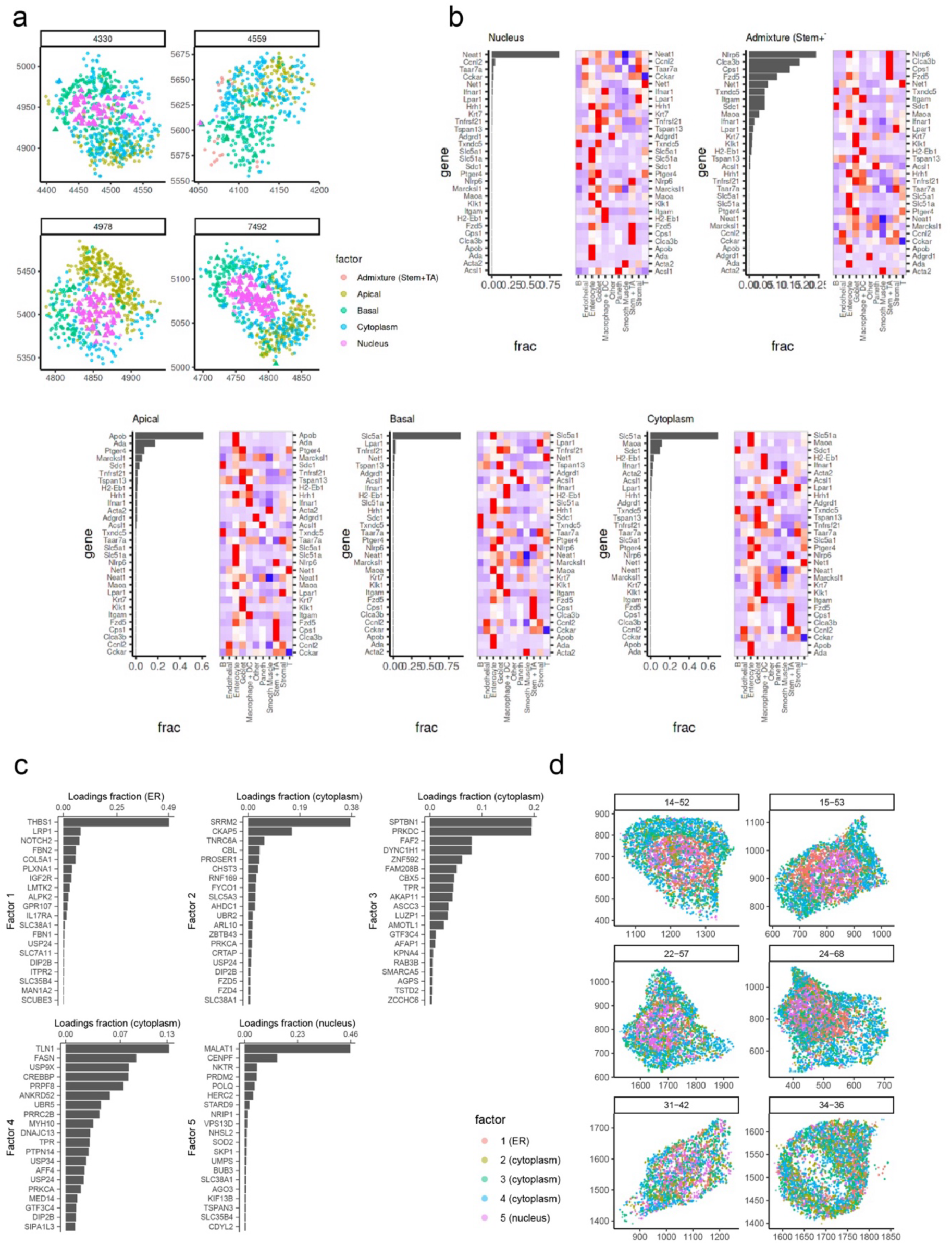
NMF factors can also extract patterns of biological intracellular variation. a. Visualization of subcellular compartments within enterocyte cells from the mouse ileum dataset. Colors indicate molecules assigned to different compartments / Factors by CRF. b. Top NMF factor loadings for NMF run on enterocytes. For each NMF factor, the left panel shows factor loadings on genes. The right panel shows a heatmap of normalized gene expression (z-scores) in different cell types, where blue indicates low expression and red indicates high expression. c. Top NMF factor loadings for NMF run on US-O2 osteosarcoma cell line dataset. For each NMF factor, the left panel shows factor loadings on genes. d. Visualization of subcellular compartments within US-O2 osteosarcoma cells.

## References

1. Palla, G., Fischer, D. S., Regev, A. & Theis, F. J. Spatial components of molecular tissue biology. Nat Biotechnol 40, 308–318 (2022).

2. Armingol, E., Officer, A., Harismendy, O. & Lewis, N. E. Deciphering cell–cell interactions and communication from gene expression. Nat Rev Genet 22, 71–88 (2021).

3. Fang, S. et al. Computational Approaches and Challenges in Spatial Transcriptomics. Genomics Proteomics Bioinformatics 21, 24–47 (2023).

4. Wu, L. et al. Spatially-resolved transcriptomics analyses of invasive fronts in solid tumors. 2021.10.21.465135 Preprint at 10.1101/2021.10.21.465135 (2021).

5. Liu, S. et al. Spatially mapping T cell receptors and transcriptomes reveals distinct immune niches and interactions underlying the adaptive immune response. Immunity 55, 1940–1952.e5 (2022).

6. Avraham-Davidi, I. et al. Integrative single cell and spatial transcriptomics of colorectal cancer reveals multicellular functional units that support tumor progression. 2022.10.02.508492 Preprint at 10.1101/2022.10.02.508492 (2022).

7. Zhao, T. et al. Spatial genomics enables multi-modal study of clonal heterogeneity in tissues. Nature 601, 85–91 (2022).

8. Toninelli, M., Rossetti, G. & Pagani, M. Charting the tumor microenvironment with spatial profiling technologies. Trends in Cancer 9, 1085–1096 (2023).

9. Hirz, T. et al. Dissecting the immune suppressive human prostate tumor microenvironment via integrated single-cell and spatial transcriptomic analyses. Nat Commun 14, 663 (2023).

10. Janesick, A. et al. High resolution mapping of the tumor microenvironment using integrated single-cell, spatial and in situ analysis. Nat Commun 14, 8353 (2023).

11. Chen, J. H. et al. Human lung cancer harbors spatially organized stem-immunity hubs associated with response to immunotherapy. Nat Immunol 25, 644–658 (2024).

12. Allen, W. E., Blosser, T. R., Sullivan, Z. A., Dulac, C. & Zhuang, X. Molecular and spatial signatures of mouse brain aging at single-cell resolution. Cell 186, 194–208.e18 (2023).

13. Kilfeather, P., et al. Single-cell spatial transcriptomic and translatomic profiling of dopaminergic neurons in health, aging, and disease. Cell Reports 43, (2024).

14. Stickels, R. R. et al. Highly sensitive spatial transcriptomics at near-cellular resolution with Slide-seqV2. Nat Biotechnol 39, 313–319 (2021).

15. Chen, A. et al. Spatiotemporal transcriptomic atlas of mouse organogenesis using DNA nanoball-patterned arrays. Cell 185, 1777–1792.e21 (2022).

16. Farah, E. N. et al. Spatially organized cellular communities form the developing human heart. Nature 627, 854–864 (2024).

17. Cable, D. M. et al. Robust decomposition of cell type mixtures in spatial transcriptomics. Nat Biotechnol 40, 517–526 (2022).

18. Moffitt, J. R. et al. High-throughput single-cell gene-expression profiling with multiplexed error-robust fluorescence in situ hybridization. Proc Natl Acad Sci U S A 113, 11046–11051 (2016).

19. Wang, X. et al. Three-dimensional intact-tissue sequencing of single-cell transcriptional states. Science 361, eaat5691 (2018).

20. Stringer, C., Wang, T., Michaelos, M. & Pachitariu, M. Cellpose: a generalist algorithm for cellular segmentation. Nat Methods 18, 100–106 (2021).

21. Greenwald, N. F. et al. Whole-cell segmentation of tissue images with human-level performance using large-scale data annotation and deep learning. Nat Biotechnol 40, 555– 565 (2022).

22. Petukhov, V. et al. Cell segmentation in imaging-based spatial transcriptomics. Nat Biotechnol 40, 345–354 (2022).

23. Park, J. et al. Cell segmentation-free inference of cell types from in situ transcriptomics data. Nat Commun 12, 3545 (2021).

24. Moffitt, J. R. et al. Molecular, Spatial and Functional Single-Cell Profiling of the Hypothalamic Preoptic Region. Science 362, eaau5324 (2018).

25. He, S. et al. High-plex imaging of RNA and proteins at subcellular resolution in fixed tissue by spatial molecular imaging. Nat Biotechnol 40, 1794–1806 (2022).

26. Arnol, D., Schapiro, D., Bodenmiller, B., Saez-Rodriguez, J. & Stegle, O. Modeling Cell-Cell Interactions from Spatial Molecular Data with Spatial Variance Component Analysis. Cell Rep 29, 202–211.e6 (2019).

27. Dries, R. et al. Giotto: a toolbox for integrative analysis and visualization of spatial expression data. Genome Biology 22, 78 (2021).

28. Garcia-Alonso, L. et al. Mapping the temporal and spatial dynamics of the human endometrium in vivo and in vitro. Nat Genet 53, 1698–1711 (2021).

29. Tanevski, J., Flores, R. O. R., Gabor, A., Schapiro, D. & Saez-Rodriguez, J. Explainable multiview framework for dissecting spatial relationships from highly multiplexed data. Genome Biology 23, 97 (2022).

30. Jerby-Arnon, L. & Regev, A. DIALOGUE maps multicellular programs in tissue from single-cell or spatial transcriptomics data. Nat Biotechnol 40, 1467–1477 (2022).

31. Shao, X. et al. Knowledge-graph-based cell-cell communication inference for spatially resolved transcriptomic data with SpaTalk. Nat Commun 13, 4429 (2022).

32. Cang, Z. et al. Screening cell–cell communication in spatial transcriptomics via collective optimal transport. Nat Methods 20, 218–228 (2023).

33. Pham, D. et al. Robust mapping of spatiotemporal trajectories and cell–cell interactions in healthy and diseased tissues. Nat Commun 14, 7739 (2023).

34. Fischer, D. S., Schaar, A. C. & Theis, F. J. Modeling intercellular communication in tissues using spatial graphs of cells. Nat Biotechnol 41, 332–336 (2023).

35. Mason, K. et al. Niche-DE: niche-differential gene expression analysis in spatial transcriptomics data identifies context-dependent cell-cell interactions. Genome Biology 25, 14 (2024).

36. Mitchel, J. et al. Tensor decomposition reveals coordinated multicellular patterns of transcriptional variation that distinguish and stratify disease individuals. 2022.02.16.480703 Preprint at 10.1101/2022.02.16.480703 (2022).

37. Barkas, N. et al. Joint analysis of heterogeneous single-cell RNA-seq dataset collections. Nature Methods 16, 695–698 (2019).

38. Wang, Y., et al. iTALK: An R Package to Characterize and Illustrate Intercellular Communication. 507871 https://www.biorxiv.org/content/10.1101/507871v1 (2019) doi:10.1101/507871.

39. Zilionis, R. et al. Single cell transcriptomics of human and mouse lung cancers reveals conserved myeloid populations across individuals and species. Immunity 50, 1317–1334.e10 (2019).

40. Haber, A. L. et al. A single-cell survey of the small intestinal epithelium. Nature 551, 333– 339 (2017).

41. Paszke, A., et al. PyTorch: An Imperative Style, High-Performance Deep Learning Library. Preprint at 10.48550/arXiv.1912.01703 (2019).

42. Kingma, D. P. & Ba, J. Adam: A Method for Stochastic Optimization. Preprint at 10.48550/arXiv.1412.6980 (2017).

43. Gaujoux, R. & Seoighe, C. A flexible R package for nonnegative matrix factorization. BMC Bioinformatics 11, 367 (2010).

44. Wang, G., Kossenkov, A. V. & Ochs, M. F. LS-NMF: A modified non-negative matrix factorization algorithm utilizing uncertainty estimates. BMC Bioinformatics 7, 175 (2006).

